# Combinatorial recognition of clustered RNA elements by a multidomain RNA-binding protein, IMP3

**DOI:** 10.1101/398008

**Authors:** Tim Schneider, Lee-Hsueh Hung, Masood Aziz, Anna Wilmen, Stephanie Thaum, Jacqueline Wagner, Robert Janowski, Simon Müller, Stefan Hüttelmaier, Dierk Niessing, Michael Sattler, Andreas Schlundt, Albrecht Bindereif

## Abstract

How multidomain RNA-binding proteins recognize their specific target sequences, based on a combinatorial code, represents a fundamental unsolved question and has not been studied systematically so far. Here we focus on a prototypical multidomain RNA-binding protein, IMP3 (also called IGF2BP3), which contains six RNA-binding domains (RBDs): four KH and two RRM domains. We have established an integrative systematic strategy, combining single-domain-resolved SELEX-seq, motif-spacing analyses, *in vivo* iCLIP, functional validation assays, and structural biology. This approach identifies the RNA-binding specificity and RNP topology of IMP3, involving all six RBDs and a cluster of up to five distinct and appropriately spaced CA-rich and GGC-core RNA elements, covering a >100 nucleotide-long target RNA region. Our generally applicable approach explains both specificity and flexibility of IMP3-RNA recognition, providing a paradigm for the function of multivalent interactions with multidomain RNA-binding proteins in gene regulation.

## Introduction

The insulin-like growth factor 2 mRNA binding protein 3 (IMP3 or IGF2BP3) belongs to a family of three highly conserved RNA-binding proteins (IMP1, IMP2 and IMP3) that are involved in post-transcriptional gene regulation of mRNAs^1^. The three mammalian paralogs are often described as oncofetal due to their expression primarily during embryogenesis and severe phenotypes in case of impaired expression^2,3^.

The currently best-understood IMP-mediated mechanism of modulating mRNA fate comprises the so-called safe-housing of specific transcripts in mRNP granules^4^. This “caging” of mRNAs ranges in its functional spectrum from packaging for cytoplasmic transport^5^, delayed translation within stable mRNPs^6-8^, cytoplasmic storage, and protection against pre-mature miRNA-directed mRNA regulation^3,9-12^. Several target mRNAs have been suggested^3,13^, with IMP1 associating with the *ACTB* mRNA zipcode element and all three IMPs regulating *HMGA2* stability via the 3´-UTR as the currently best-studied examples^9-12,14-16^.

In contrast to IMP1 and IMP2, the biological relevance of IMP3 has long been underestimated. Research on IMP3 largely focused on its association with many cancer-related tumor entities, since its re-expression correlates with a poor prognosis for patients, classifying IMP3 as a tumor marker^17-19^.

The IMP protein family represents a prototypical example of multidomain-RBPs and is characterized by a common architecture of six potential RNA-binding units: two N-terminal RNA-recognition motifs (RRMs) and four consecutive hnRNP K-homology (KH) domains^1^. It has been a longstanding question how multiple RBDs cooperate in specific and high-affinity RNA-target recognition: Which of the individual domains are involved, what are their contributions, and how flexible is the RNA-protein interaction pattern?

Assessing the contributions and cooperativity of multiple RBDs in binding to multipartite RNA motifs is challenging, and a generally applicable approach has not been described so far. Due to the potential dynamic domain arrangements of multiple RBDs, structural studies require an integrated approach, combining solution techniques and crystallography^20-24^. For the IMPs, structural information is available only for single RRMs of IMP2 (RRM1, PDB-ID: 2CQH) and IMP3 (RRM2, PDB-ID: 2E44, both unpublished). The presence of a very short linker sequence suggests that the two domains are arranged in a compact tandem, which might drive their RNA specificity. Analogously, there is evidence that the KH1-2 and KH3-4 tandem domains represent pre-arranged RNA-binding modules for recognition of bipartite RNA sequence motifs. Structures of the human IMP1 KH3-4^14^, as well as the KH3-4 di-domains of the chicken ortholog ZBP1^16^ proved the existence of an extended domain interface between KH3 and 4. These structures suggest target RNA motifs to require a minimal spacing to be recognized by the tandem RBDs. For example, KH3-4 of ZBP1/IMP1 recognizes a combination of two sequence elements: CGGAC-N_10-25_-(C/A-CA-C/U) in both possible arrangements^14-16^.

Previous studies proposed short recognition sequences of IMPs, based on *in vivo* CLIP^3,13,25^ and *in vitro* selections (SELEX, RNAcompete and Bind-N-seq)^5,26-28^, all suggesting an overall CA-rich consensus. However, the major limitation of *in vitro* selection approaches is that they usually start with short degenerate sequences, which can accommodate only a single RNA-binding motif. Therefore the contributions of individual domains have remained elusive. Finally, while previous studies provide evidence for an essential role for KH domains in RNA interaction, no function had been ascribed yet to the two RRMs^5,14-16,29,30^.

To study IMP3 as a prototypical example of a multidomain-RBP we established a systematic, domain-resolved SELEX procedure coupled with RNA-seq and combinatorial bioinformatic approaches. Importantly, we used a very long degenerate sequence (N_40_) as a basis for SELEX, to allow multiple RNA contacts with more than a single RNA-binding domain, and a corresponding bioinformatic spacing analysis. This led us to the discovery that IMP3 recognizes – through the activity of all of its six RNA-binding domains-an extended array of multiple *cis*-acting RNA elements, comprised of CA-rich motifs and sequences with a common GGC core. These biochemical findings are supported by integrated structural biology, combining crystallography and NMR for structural analysis and RNA-binding studies of IMP3 KH and RRM-tandem domains.

Taken together, we provide biochemical, bioinformatic, and structural evidence for recognition of an ordered array of RNA elements by IMP3, arranged in a certain spacing pattern and covering regions that can span more than 100 nts. This model is supported by the analysis of endogenous IMP3 target mRNAs, including the well-studied *HMGA2* transcript, for which we investigated the functional cross-regulation between IMP3 and the let-7 miRNA. In sum, we provide a new framework for investigating large regulatory mRNP complexes. Thereby, we have established a general approach to systematically dissect complex and combinatorial RNP networks, which can be applied to any multidomain RNA-binding protein.

## Results

### IMP3 recognizes an array of distinct sequence elements: identification by SELEX-seq

To dissect the complex RNA-binding properties of IMP3, we used individual, GST-tagged subdomains and applied an *in vitro* SELEX procedure, including four rounds of selection with a random N_40_-RNA pool and subsequent RNA-seq analysis (**Fig. 1a, b,** and **Supplementary Fig.1**). Note that instead of standard short degenerate regions, we used an N_40_-RNA pool to be able to dissect and analyze arrays of several motifs, including their spacing; in addition, we sequenced after each round of selection, which allowed monitoring sequence enrichment throughout the SELEX procedure.

**Figure 1.**
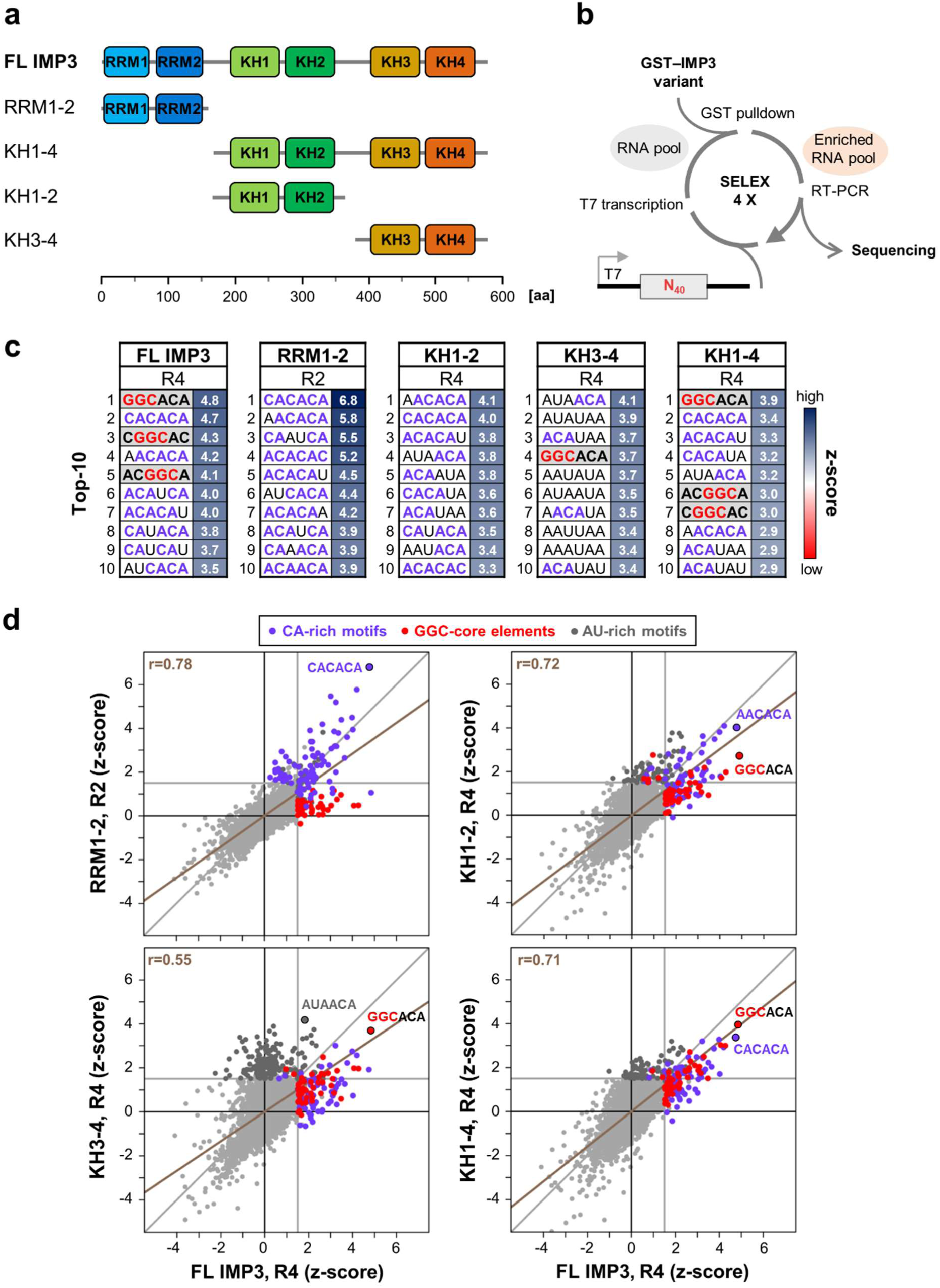
SELEX-seq analysis of IMP3 RNA-binding motifs. (**a**) Truncated IMP3 derivatives that were used for SELEX experiments (FL = full-length). RNA-binding domains are color-coded. (**b**) SELEX-seq procedure. Using GST-tagged IMP3 truncations (GST alone as negative and full-length IMP3 as positive control) and an N_40_-RNA pool, sequences bound by respective proteins were enriched through four SELEX rounds and analyzed by sequencing after each round. (**c**) Top-10 enriched 6-mer motifs for all IMP3 derivatives measured by z-score after the fourth round of selection (R4), except for RRM1-2 (R2, for the complete dataset, see **Supplementary Table 1**). CA-rich motifs are highlighted in violet, elements with a common GGC consensus in red with grey background. (**d**) Correlation of 6-mer motif enrichment (measured by z-score) for IMP3 truncations (y-axis) in comparison to the positive control, full-length IMP3 (x-axis). Motifs with z-scores higher than 1.5 (vertical/horizontal grey lines) in either x-or y-axis are highlighted in violet for CA-rich motifs, red for GGC-core elements, and dark grey for AU-rich motifs. Pearson´s correlation by linear regression is shown as a brown line with correlation coefficients (r) indicated.

Single domains, such as RRM1 or KH1, did not show RNA-binding activity (data not shown). In addition, previous structural studies had shown that at least the KH domains 3-4 of the related ZBP1/IMP1 are organized as a functional pseudo-dimer (see *Introduction*). Therefore, we relied on truncated tandem domains for our analyses: RRM1-2, KH1-2, KH3-4, as well as an extended version containing all four KH-domains, KH1-4 (**Fig. 1a** and **Supplementary Fig. 1**). In parallel, full-length IMP3 (as positive control) and GST alone (as negative control and for background correction) were analyzed. Motif-enrichment analysis by z-score calculation was performed for all possible 4-, 5-and 6-mers, and were corrected at each round with the corresponding GST SELEX round (top-10 enriched 6-mer motifs in **Fig. 1c**; complete dataset in **Supplementary Table 1**). In parallel, the correlation of motif-enrichment datasets was tested for each tandem domain by comparison with the positive control, full-length IMP3 (**Fig. 1d**).

For the full-length IMP3 protein, this SELEX analysis revealed two populations of enriched motifs, CA-rich motifs as well as motifs with a GGC core (<u>GGC</u>A and C<u>GGC</u>; **Fig. 1c**). The KH1-4 variant, which lacks the N-terminal RRM domains, showed a very similar motif enrichment as the full-length protein, revealing that the four KH domains recognize both types of motifs (**Fig. 1c,d**). Separate analysis of KH1-2 and KH3-4 tandem domains also showed the enrichment of GGC-core elements within the top-30 hexamers (**Supplementary Table 1**), but the most-enriched sequences were either CA-(KH1-2) or CA/AU-rich (KH3-4), indicating that at least one of the KH domains of each tandem binds such a sequence (**Fig. 1c,d**, for the enrichment of AU-sequences, in particular by KH3-4, see *Discussion*).

Most surprisingly, we found that RRM1-2, which until now had been described as non-functional in RNA binding, in fact exhibited a high preference for CA-rich and CA-repeat sequences, but not for the GGC-core elements (**Fig. 1c,d**). This specificity was observed after the second SELEX round, but was lost with more stringent washing conditions within rounds three and four. Therefore, only the first two SELEX rounds were analyzed for the RRM1-2 derivative (see *Discussion*). Furthermore, comparison of all SELEX rounds between the complete set revealed that, as expected, KH1-2, KH3-4 and the longer KH1-4 variant overlap most, whereas RRM1-2 showed the least overlap with the isolated KH domains (**Supplementary Figs. 1** and **2**).

Taken together, our findings strongly argue for differential recognition of an extended array of two different types of motifs (CA-rich and GGC-core elements), which are bound by the KH tandem domains. Besides that, we provide evidence that the RRM1-2 domains contribute additional binding of a CA-rich element.

### Spacing analysis of binding motifs: a model for RNA recognition by IMP3

To identify how the different domains of IMP3 recognize consecutive elements on a single RNA, we analyzed our SELEX-seq data for spacing between enriched 4-mer motif combinations, using a window of 0-25 nts (**Fig. 2a**). Enriched combinations of two types of motifs (CA-rich and GGC-core elements) and their spacing were measured by z-score analysis (see **Supplementary Table 2** and *Methods*).

**Figure 2.**
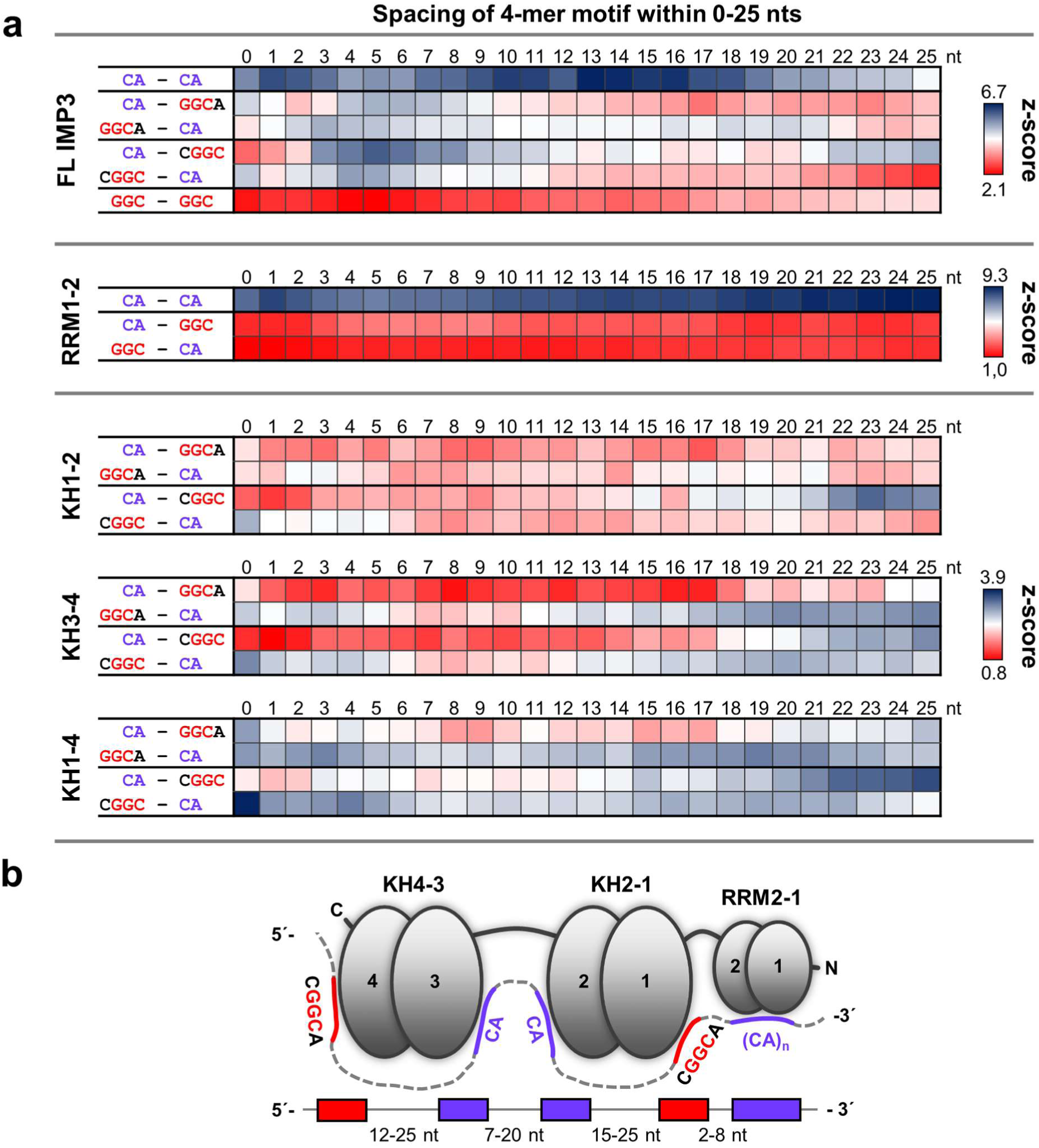
Spacing analysis reveals consensus array of IMP3-binding motifs. (**a**) Enrichment of motif combinations with spacing between 0-25 nts for the full-length IMP3 (top), and RRM1-2 (middle), KH1-2, KH3-4 and KH1-4 domains (bottom), measured by z-score and shown as heat map. The combinations of the two GGC-core elements (<u>GGC</u>A / C<u>GGC</u>) with CA-rich motifs are shown for full-length IMP3 and the KH-containing derivatives, the combinations of two GGC-core elements (GGC / GGC) for full-length IMP3 only. Spacing between CA-rich motifs was analyzed for full-length IMP3 as well as RRM1-2 (for a summary of all combinations of CA-rich and GGC-core motifs, see **Supplementary Table 2** and *Methods*). Individual z-score scales are given on the right. (**b**) Model for RNA recognition by IMP3, based on SELEX-seq analysis.

Analysis of the full-length IMP3 data showed that the most enriched motif combinations were either two CA-rich motifs with a short or medium-range spacing (CA-N_0-3_-CA; CA-N_7-20_-CA; with a maximum at N_13-16_), or a combination of a CA-rich motif with one of the identified GGC-core elements. For all combinations (CA-<u>GGC</u>A, <u>GGC</u>A-CA, CA-C<u>GGC</u> and C<u>GGC</u>-CA) we observed shorter spacing of N_2-11_ nucleotides, with a maximum at N_4-6_. However, longer spacing was found to be clearly specific for either one of the two very similar GGC-elements (<u>GGC</u>A versus C<u>GGC</u>): Only <u>GGC</u>A-N_16-21_-CA or CA-N_21-25_-C<u>GGC</u> were enriched, but not the respective reverse orientations (**Fig. 2a**, top). This indicates that, first, these sequence elements need to be appropriately spaced for recognition by IMP3; second, the arrangement of two motifs relative to each other is essential, and third, that both GGC-core elements seem to be differentially recognized. Finally, combinations of two GGC elements were not enriched.

Next, we applied this approach to the KH subdomains to obtain a refined view of motif spacing for IMP3. For each of the KH1-2, KH3-4 and KH1-4 subdomains we analyzed spacing between either one of the two GGC-core elements (<u>GGC</u>A versus C<u>GGC</u>): and the respective combination with CA-rich motifs identified through analysis of the full-length protein (**Fig. 2a**, bottom).

Strikingly, we found that the KH1-2 subdomain shows a preference only for the combination of CA-rich motifs and the C<u>GGC</u>-element in one of the possible orientations, with a strong CA-N_17-25_-C<u>GGC</u> spacing optimum. At the same time, we observed selection against the three other combinations, underlining high specificity for both the relative arrangement of CA and GGC motifs, as well as for one type of GGC-core element (C<u>GGC</u>). This observation is supported by the results obtained for the full-length IMP3 protein (**Fig. 2a**, top).

In contrast, KH3-4 showed the strongest enrichment for <u>GGC</u>A-N_12-25_-CA, but – to a similar extent – appears to tolerate also C<u>GGC</u> in combination with a CA-rich motif, in either orientation and with a spacing of N_18-25_ and N_12-25_, respectively. Similar to full-length IMP3 and KH1-2, the CA-<u>GGC</u>A motif combination was found to be least enriched for KH3-4.

Finally, for KH1-4, we detected a mix of enriched motif spacing already observed for the separate KH1-2 and KH3-4 domains, with a preference for both <u>GGC</u>A-N_12-25_-CA and CA-N_15-25_-C<u>GGC</u> orientations, tolerating also C<u>GGC</u>-N_12-25_-CA (**Fig. 2a**, bottom; see *Discussion*).

In addition, spacing analysis for RRM1-2 revealed strong enrichment for CA-rich motif combinations in all positions within the 25 nts window, but not for the GGC-core elements (**Fig. 2a**, middle), again arguing for a high preference for extended CA-rich repeat elements, in agreement with our previous analyses (**Fig. 1c,d**, see *Discussion*).

Based on these datasets, we assembled a model how IMP3 recognizes RNA (**Fig. 2b**). Due to the selective enrichment of specific motif arrangements and the known sequence preference of KH3-4 subdomains of the IMP1 paralog (see *Introduction*), we propose that KH1 and KH4 each recognize sequence elements with a common GGC core, whereas KH2 and KH3 bind to CA-rich motifs. The RRMs may provide an additional, stabilizing interaction with adjacent CA-rich motifs. It should be noted, that due to the symmetry of this array of sequence elements, our spacing analysis would partially support both polarities of IMP3 binding to its target RNAs.

### *In vitro* analysis of IMP3 RNA recognition

To validate our model presented in **Fig. 2b**, we designed an RNA sequence based on our SELEX analysis, containing domain-specific minimal 4-mer sequence elements that are appropriately spaced by unrelated sequences, extending to a total length of 101 nts (101-mer RNA): <u>GGC</u>A-N_20_-CACA-N_14_-CACA-N_22_-C<u>GGC</u>-N_4_-(CA)_4_; (**Fig. 3a**, for the full sequence, see **Fig. 7a**).

**Figure 3.**
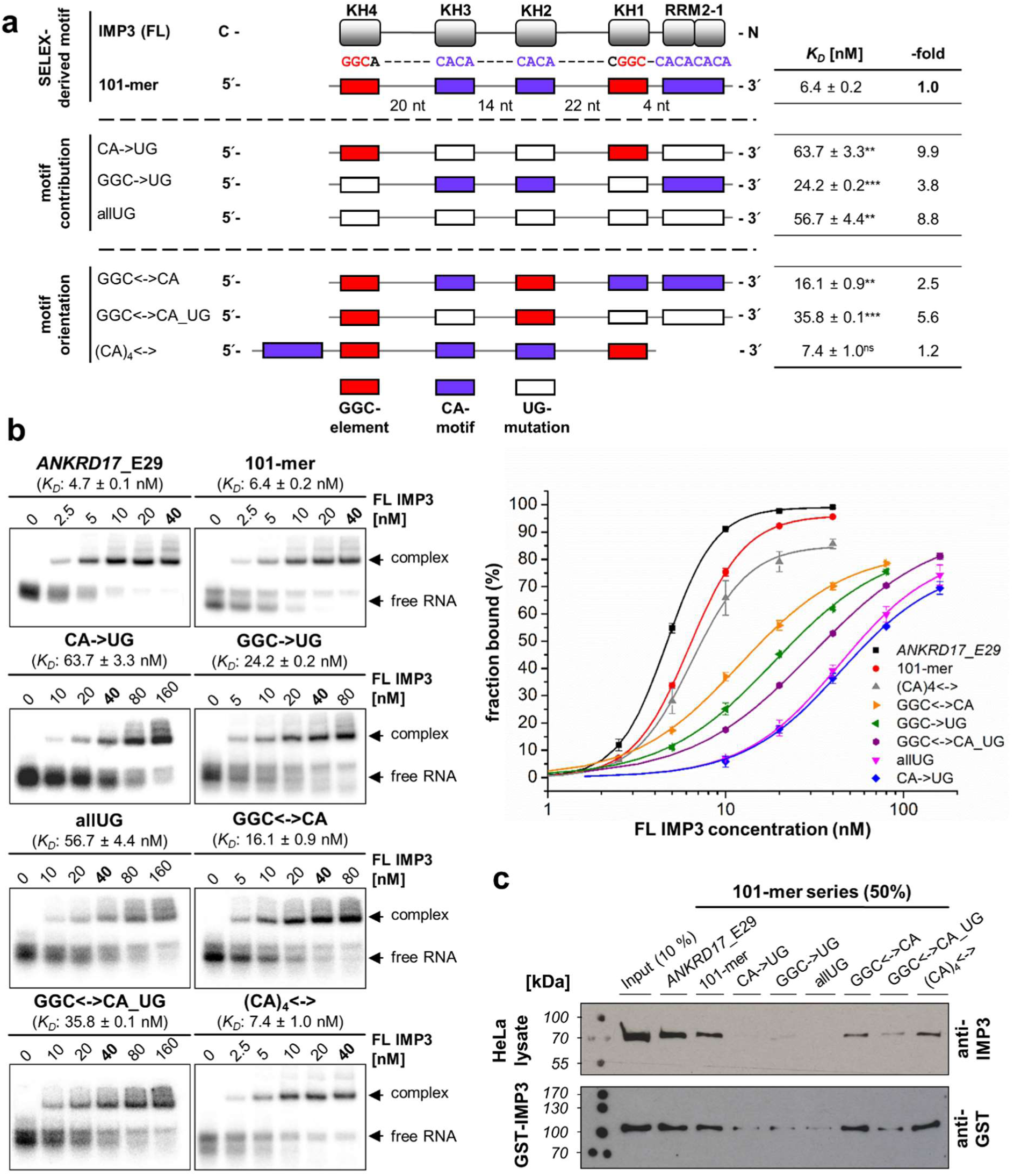
Validation of the SELEX-derived array of IMP3-binding motifs: mutational analysis. (**a**) Design of a 101-mer RNA, containing all SELEX-derived IMP3-binding motifs (GGC-motifs, red boxes; CA-motifs, violet boxes) with appropriate spacing and serving as a basis for mutational analysis and validation assays. The IMP3 domains potentially interacting with respective sequence elements of the 101-mer RNA are indicated (top). The contributions of specific motifs were tested by mutational analysis (CA-motifs or GGC-core elements or both of them mutated to UG; middle). The importance of motif orientation was analyzed by shuffling of domain-specific sequence motifs (KH1-2-specific motifs: GGC<->CA; additional substitution of the CA-motifs: GGC<->CA_UG; relative positioning of the RRM1-2-specific motif: (CA)_4_<->; bottom). *K*_D_ values obtained by electromobility shift assays (EMSAs, see panel **b**) and respective changes in binding affinity (-fold) compared to the wildtype 101-mer sequence are summarized on the right (p<0.005^**^, p<0.001^***^, two-sided t-test). (**b**) IMP3 interaction with RNAs of the 101-mer series, assayed by EMSAs. Full-length protein (0-40, 0-80 or 0-160 nM) was titrated to a constant concentration of respective ^32^P-labeled RNAs (5 nM). A 121-nt region from the IMP3 target mRNA *ANKRD17* (exon 29) served as a positive control. Corresponding binding curves for *K*_D_-estimation are shown on the right (mean and standard deviation of three experiments). (**c**) Pulldown of endogenous IMP3 in HeLa cell lysate (top) or of recombinant GST-IMP3 (bottom) with 3´-biotinylated RNAs of the 101-mer series. IMP3 was detected by Western blot with either IMP3-(top) or GST-specific antibodies (bottom).

The 101-mer RNA was used as a basis for mutational analysis to determine the contribution of individual sequence elements to the overall affinity of the protein. Electromobility shift assays (EMSAs) revealed that the full-length protein recognizes the ^32^P-labeled 101-mer RNA with high affinity (dissociation constant *K*_D_ = 6.4 ± 0.2 nM, **Fig. 3a,b**), comparable to the positive control, a sequence of similar length derived from exon 29 of the *ANKRD17* transcript (121 nts, *K*_D_ = 4.7 ± 0.1 nM, **Fig. 3a,b**). The *ANKRD17* transcript had been recently identified by us as strongly IMP3-associated^27^ and harboring nearly the exact array of sequence elements proposed in our 101-mer (see also **Fig. 7a**).

To test for motif contribution within the 101-mer sequence, we either substituted the CA-motifs (CA->UG), the GGC-core elements (GGC->UG), or a combination of both (allUG), each by mutating to UG (for full sequences, see **Supplementary Table 3**). Substitution of the GGC-core elements led to a 4-fold reduction in affinity, and mutation of the CA-motifs, or the combination of both, led to a 9-to 10-fold reduction (**Fig. 3a,b**). This indicates that both elements are important for high-affinity RNA recognition.

We also evaluated the importance of motif orientation, by changing the order of KH1-2-specific elements (GGC<->CA), resulting in a 2.5-fold decrease in affinity (**Fig. 3a,b**). The additional substitution of CA-motifs within this context (GGC<->CA_UG) led to an further reduction (almost 6-fold total). This shows that the protein prefers the SELEX-derived orientation of elements, but can adapt to changes with relatively modest effects on binding affinity. Furthermore, we tested the influence of the CA-repeat element, which is located on the very 3´-end and recognized by RRM1-2, by moving it to the 5´-end ((CA)_4_<->). Surprisingly, binding affinity remained unchanged, suggesting that either this element does not significantly contribute to overall affinity or that IMP3 can recognize the element in both positions, consistent with our spacing analysis (see **Fig. 2**).

Our EMSA-based results were consistent with pulldown assays of endogenous IMP3 protein from HeLa cell lysate as well as of recombinant GST-tagged IMP3 with 3´-biotinylated RNAs and subsequent Western blot detection (**Fig. 3c**).

In sum, these consistent results from biochemical assays, quantitative EMSA and semi-quantitative pulldown strongly support our proposed model of target RNA recognition involving all IMP3 RBDs (**Fig. 2b**).

### Structure and RNA recognition by the IMP3 tandem KH1-2 domain

Given substantial primary sequence conservation of the IMP1 and IMP3 KH3-4 tandem domains (**Supplementary Fig. 3**), similar RNA-binding features were expected for IMP3 KH3-4, as suggested by Chao and colleagues^14^. In contrast, the RNA recognition by the IMP3 KH1-2 tandem had so far not been analyzed. To determine the individual contributions of KH1 and 2 (Lys192 to Ile355), their RNA binding was inactivated by mutation (GKEG motif to GDDG), while maintaining the crucial tandem context^14-16^, resulting in four possible combinations (**Fig. 4a**). Our NMR data clearly proved the integrity of all constructs (**Supplementary Fig. 3**). We analyzed crystals of both wildtype KH1-2 and KH1-Δ2 versions for structural characterization. While the former only generated very low-resolution diffraction data, we were able to solve the structure of KH1-Δ2 at 2.0 Å resolution (**Fig. 4b** and **Supplementary Table 4**). SAXS (small angle X-ray scattering) data back-calculated based on the crystal structure are in good agreement, indicating that the crystal structure reflects the monomeric solution geometry (**Fig. 4c**), which also closely resembles other tandem KH domains (**Supplementary Fig. 3**). We conclude that the IMP3 KH1-2 tandem is a stable monomeric folding unit.

**Figure 4.**
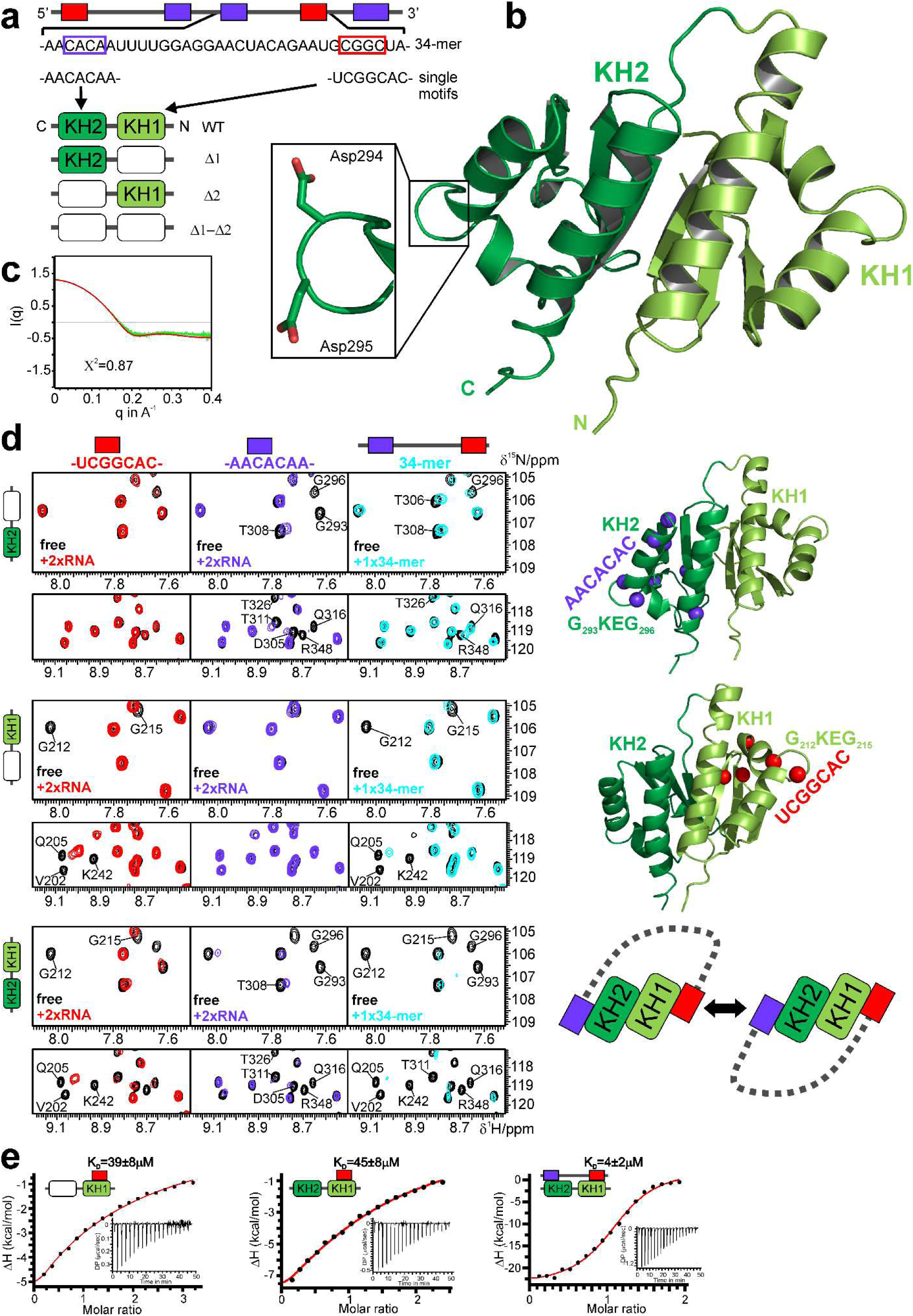
Structure and RNA recognition of the IMP3 tandem KH1-2 domain. (**a**) Protein constructs and RNAs used. (Top) Scheme of the 101-mer RNA region, which includes the 34-mer sequence (below), covering the cognate binding region of the KH1-2 domain. The two recognition sequences for KH1 and KH2 are embedded in two respective 7-mers. (Bottom) Wildtype (WT) and three different versions of KH1-2 (in Δ-versions of the domains, GKEG replaced by GDDG^30^). A proof of concept for this approach is shown in **Supplementary Fig. 3**. (**b**) Crystal structure of the KH1-Δ2 tandem domain (see also **Supplementary Table 4** and **Supplementary Fig. 3**). The zoom-in shows the mutated GKEG loop with two aspartates replacing Lys294 and Glu295 in KH2. (**c**) SAXS curve of KH1-Δ2 at 4 mg/ml and overlaid with a theoretical curve from the crystal structure in **b**) created by *Crysol* (red) ^31^ (**d**) HSQC overlays showing KH1-2 versions Δ1 (upper), Δ2 (middle) and WT (lower row) free (black) and when bound to two-fold excess of either of the short RNAs or equimolar 34-mer RNA (see color code). Two different spectral regions (top/bottom) are shown. Selected residues as representative probes in the active subdomains (light/dark green color for KH1 and KH2, respectively), are annotated in the spectra. Amide groups of strongly affected residues are shown as spheres in the structures on the right. The scheme at the lower right suggests two possible modes of KH1-2 interacting with the 34-mer RNA. Complete NMR spectra and CSP plots are provided in **Supplementary Fig. 4** and **5**. (**e**) Representative ITC curves for binding of KH1 (in the KH1-Δ2 context) and KH1-2 WT when titrated with UCGGCAC. The plot on the right shows the binding of KH1-2 WT to the 34-mer RNA comprising both motifs. The suggested topology of the protein-RNA complex and dissociation constants (*K*_D_) for the interaction are indicated (mean and standard deviation of three experiments). All ITC measurements are summarized in **Supplementary Table 5**.

We next examined RNA-binding contributions of the KH1 and KH2 domains by inactivation of the individual domains in the KH1-2 context, using SELEX-derived 7-mers from the rationally designed 101-mer (**Fig. 3, 4a,d** and **Supplementary Figs. 4** and **5**). First, NMR was used to identify the RNA sequence recognized by the individual subdomains in (**Fig. 4d**). Indeed, KH1 clearly favors binding of the GGC-motif, while KH2 prefers binding to the CA-RNA. We did not see mentionable cross-reactivity of domains with the respective unrelated RNA in the context of single KH1-2 Δ-versions as shown by a full CSP analysis (**Supplementary Figs. 4** and **5**).

Can we also observe specific binding of motifs in the wildtype KH1-2 context? Here, a clear preference of KH1 for its GGC target motif was observed, while KH2 showed a lower, but significant preference for CA. Given that larger NMR CSPs were observed for the KH1/GGC, compared to the KH2/CA-RNA interaction, RNA binding appears to be mediated primarily through KH1. Indeed, ITC revealed a measurable KH1-GGC interaction in the low-to-medium micromolar range, while the KH2-CA complex could not be determined in our ITC setup (**Fig. 4e** and **Supplementary Table 5**). Notably, the respective interactions were also observed in the context of the intact wildtype KH1-2.

When both the GGC and the CA-RNA motifs are present in a single RNA ligand, an overall higher binding affinity for wildtype KH1-2 is expected. To confirm this we used a corresponding region (34-mer, **Fig. 4a**) from the 101-mer RNA, including a 22 nt linker separating the GGC-and CA-motifs, as suggested by the spacing analysis (**Fig. 2** and **3a**). As shown in **Fig. 4d**, significant CSPs were observed for KH1 and KH2 that compare well to the titration with short 7-mer GGC-and CA-RNA sequences, respectively. However, spectral changes in general appeared to be more widespread. In HSQC experiments we observed severe line broadening for most NMR signals in either subdomain upon titrating the 34-mer RNA (**Fig. 4d** and **Supplementary Fig. 3, 4** and **5**). This indicates the involvement of both domains within the RNP complex in a measurably larger and compact complex. The simultaneous recognition of both RNA motifs in a 1:1 complex requires looping of the 34-mer RNA around the KH1-2 tandem. This may occur with two possible orientations (**Fig. 4d)**, as described earlier for KH3-4 of IMP1^14,15^.

Finally, we performed ITC experiments with the wildtype KH1-2 and 34-mer RNA (**Fig. 4e** and **Supplementary Table 5**). As expected, a 10-fold higher affinity compared to the single interactions of 7-mer RNAs indicates a cooperative binding event that shifts affinity by one order of magnitude. The 1:1 stoichiometry of the KH1-2/34-mer RNA complex clearly argues for the formation of a looped-RNA-KH1-2 complex, which is also supported by a significant gain in the entropy term. Altogether, our data support the preference of KH1-2 subdomains for specific SELEX-derived RNA motifs and cooperative recognition when both motifs are present in a longer context.

### Molecular determinants of IMP3 RRM1-2-RNA interactions

To assess the RNA interactions of the IMP3 RRM1-2 domains we purified an optimized construct, which yields excellent NMR spectra, consistent with a monomeric conformation. Secondary chemical shifts reveal the presence of a canonical RRM secondary structure (**Supplementary Fig. 6**). NMR ^15^N relaxation experiments indicate a compact arrangement of domains with almost no linker flexibility, suggesting that the two domains appear as tandem (**Fig. 5a**). This is also supported by the tumbling correlation time, estimated from ^15^N *R*_1_ and *R*_2_ relaxation rates, consistent with a globular 18-kDa protein (**Fig. 5a** and **Supplementary Fig. 6**). Static-light scattering unequivocally proves the protein to be a monomer (**Supplementary Fig. 6**). SAXS data indicate a compacted arrangement of the tandem domains (**Fig. 5b)**.

**Figure 5.**
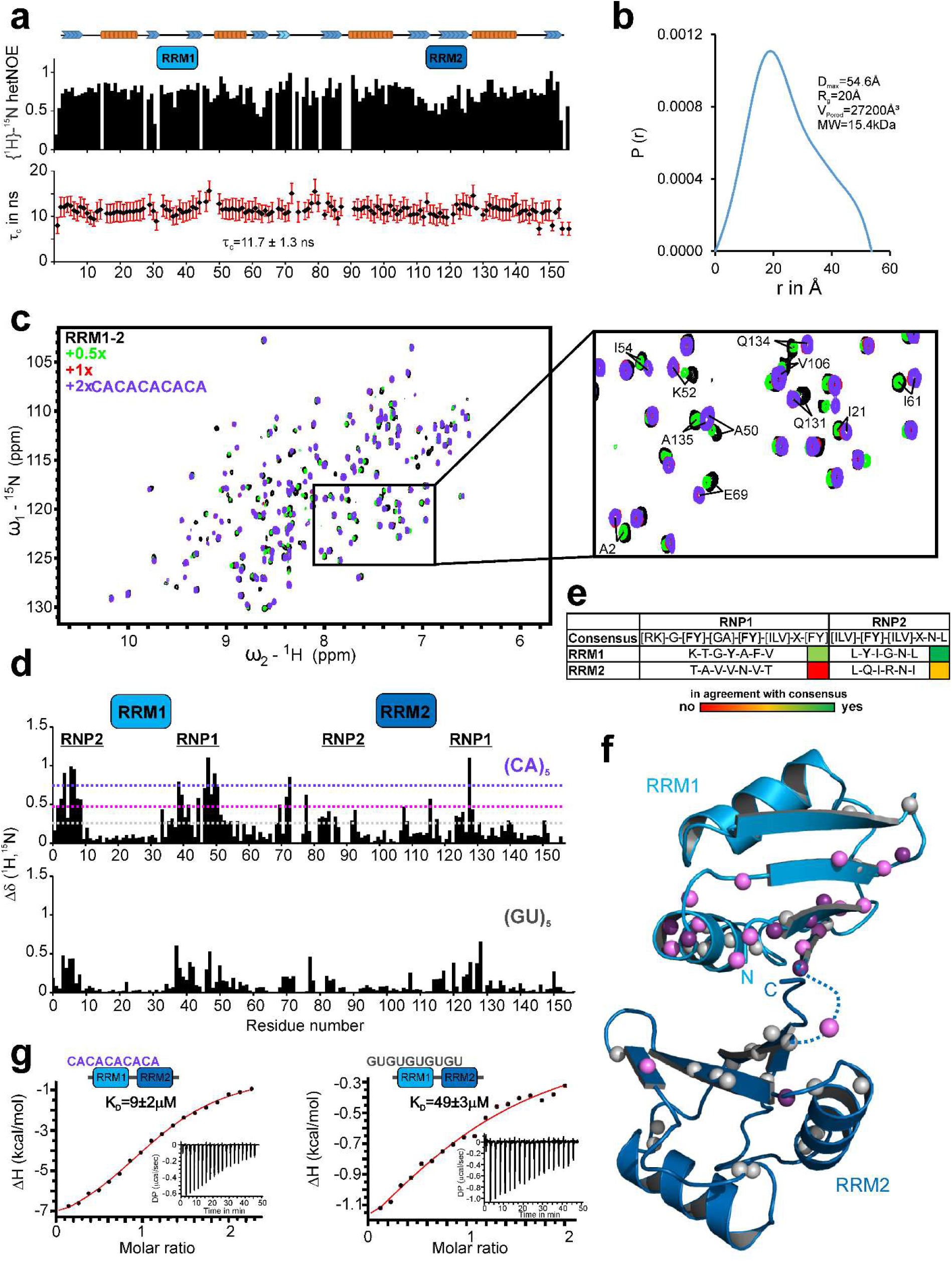
RNA recognition mode of the IMP3 RRM1-2 tandem domains. (**a**) IMP3 RRM1-2 function as tandem in solution. Secondary structure elements in the RRM1-2 tandem domains as obtained from secondary chemical shifts are shown on top. {^1^H}-^15^N heteronuclear NOE values show that the linker connecting the two globular domains is rigid. Tumbling correlation-time values (τ_C_, bottom), derived from NMR relaxation data (**Supplementary Fig. 6**), show an average value of 11.7 ns, indicating that both domains tumble together in solution. Gaps indicate prolines or residues with missing data. (**b**) Pairwise distance distribution, P(*r*), for IMP3 RRM1-2 at 1 mg/ml derived from SAXS data (**Supplementary Fig. 6**). The maximum pairwise distance (D_max_), radius of gyration (*R*_g_) and the Porod volume (V_Porod_) are consistent with a monomeric RRM1-2 tandem domain particle in solution. (**c**) Overlay of ^1^H,^15^N NMR correlation spectra of RRM1-2 alone and in presence of different concentrations of (CA)_5_ RNA (see color code). The inset shows representative residues affected by RNA binding. (**d**) Chemical shift perturbations (CSP) observed (see panel **c**) at the endpoint of the titration. The two domains and their RNP sequence motifs are labelled on top. The dotted lines indicate CSP thresholds calculated as average (grey) plus one and two standard deviations (pink and violet, respectively). The lower panel shows CSP from an NMR titration with (GU)_5_ RNA (**Supplementary Fig. 6**). (**e**) RNP sequence motifs in the RRM1 and RRM2 subdomains. (**f**) Mapping of CSPs for the titration with the (CA)_5_ RNA (panel **d**) onto a structural model of RRM1-2 (see *Results* and *Methods*). Amides are shown as spheres colored according to thresholds in panel **d**). (**g**) ITC data for the titration of RRM1-2 with (CA)_5_ or (GU)_5_ RNAs. A titration of (CA)_3_ hexamer to RRM1-2 is shown in **Supplementary Fig. 6**. The suggested complex topology and *K*_D_s are indicated. Values represent mean and standard deviation of three experiments. All ITC measurements are summarized in **Supplementary Table 5**.

We next tested binding of CA-repeat RNAs by RRM1-2 using NMR titrations. A (CA)_5_ 10-mer was chosen to potentially cover both RRMs (**Fig. 5c**). Strong NMR chemical shift perturbations were observed for residues in RRM1, while RRM2 was less affected. Hot spots map to regions around the RNP motifs (**Fig. 5d**). Interestingly, the control RNA, (GU)_5_, led to a very similar, yet much weaker pattern of CSPs in RRM1 and 2, indicating a preference for CA.

Sequence analysis suggested that RRM2 harbors a degenerate RNP2 motif and lacks a canonical RNP1 motif (**Fig. 5e**). We conclude that CSPs in RRM2 were observed because they are indirectly affected by RNA binding in RRM1 and caused by the length of the RNA. We repeated NMR titration experiments of RRM1-2 with a (CA)_3_ 6-mer RNA that should not extend towards RRM2 in the tandem domain arrangement. However, we found almost identical CSPs (**Supplementary Fig. 6**) as compared to (CA)_5_ and conclude that the two domains are arranged in a way that causes binding of RNAs through RRM1 to be sensed by nearby residues in RRM2. We derived a structural model of the RRM1-2 tandem domains filtered against SAXS data and NMR CSPs (see *Methods*)(**Fig. 5f** and **Supplementary Fig. 6**). The majority of significant CSPs localizes to RRM1, while a mentionable number of amides in RRM2 still showed CSPs above average.

Finally, ITC was used to quantify RNA binding to RRM1-2 (**Fig. 5g** and **Supplementary Table 5**). The interaction with (CA)_5_ revealed a low-micromolar affinity, and in line with our NMR data we found the same affinity for RRM1-2 when binding to the 6-mer CA-RNA (**Supplementary Fig. 6**). This supports our hypothesis where binding takes place primarily in RRM1 through an interface with not more than six nucleotides of RNA. A 5-to 6-fold lower affinity of (GU)_5_ with RRM1-2 is consistent with the reduced CSPs. However, this number still shows some non-specific RNA binding to this non-cognate motif, as often observed for canonical RRM-and KH-domains^32,33^.

In sum, we have shown that RRM1-2 significantly contributes to the overall RNA binding of IMP3 through the specific recognition of CA-rich RNAs, as suggested by our SELEX experiments.

### All tandem domains of IMP3 actively contribute to RNA recognition

To verify further the suggested concept with all IMP3 RBDs engaged in multivalent RNA recognition, we next tested the contribution of individual tandem domains within the full-length-protein context. Therefore, we mutated critical amino acids in respective domains to inactivate individual tandem domains (ΔRRM1, ΔKH1-2, ΔKH3-4 and ΔKH1-4; **Fig. 6a**), followed by EMSA assays with the designed 101-mer RNA (**Fig. 6b**). Since RRM2 does not contain well-conserved RNP motifs and consistent with our structural analysis (see above and **Fig. 5**), only RRM1 of the RRM1-2 tandem domain was mutated to assess the contribution of the RRM1-2 tandem domains. Strikingly, inactivation of RRM1 alone led to a 5-fold reduced affinity compared to wildtype (WT), indicating that this domain indeed contributes to RNA binding also in the full-length context.

**Figure 6.**
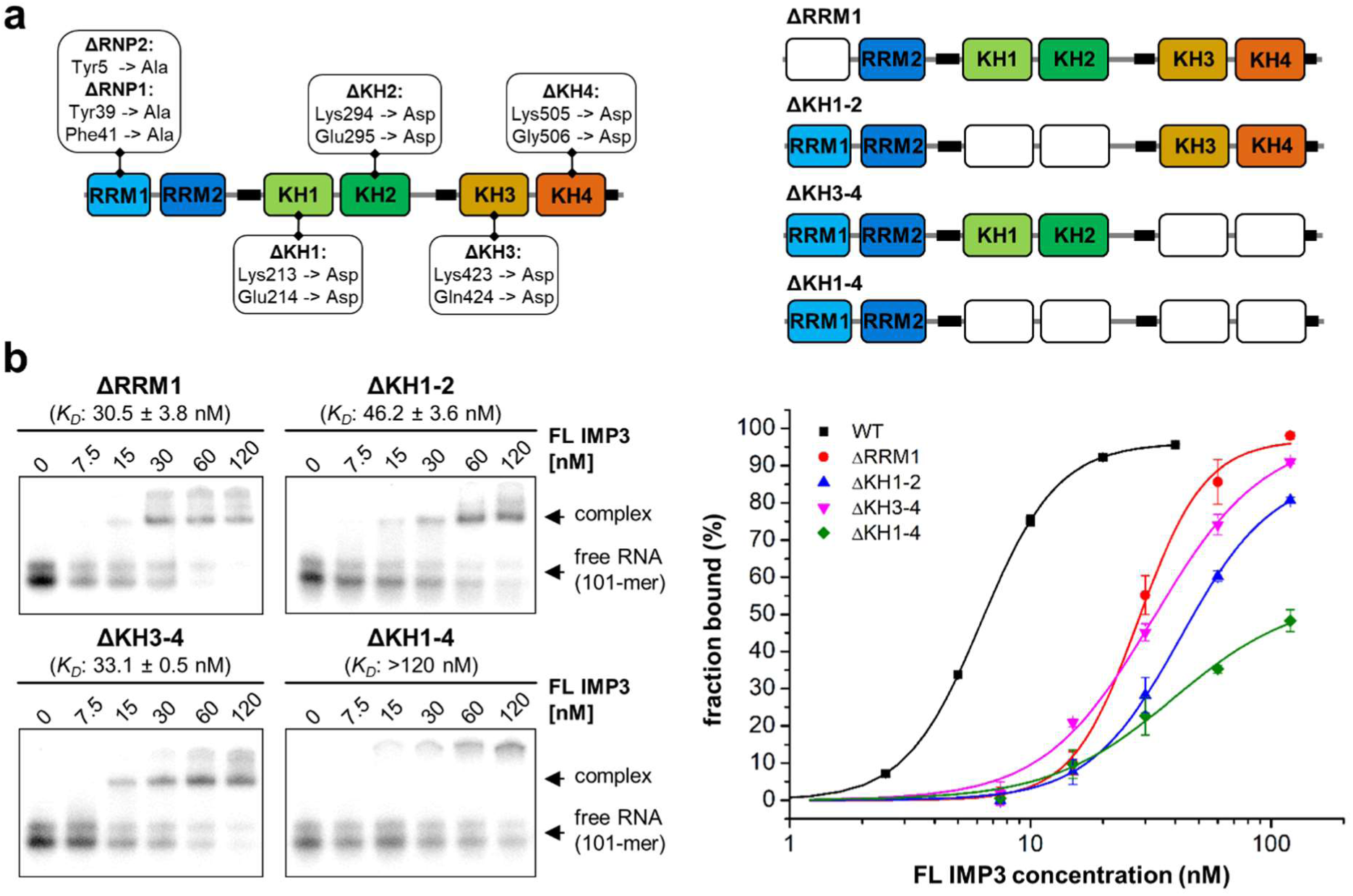
Functional analysis of individual RNA-binding domains of IMP3. (**a**) Summary of mutations introduced in full-length IMP3 for functional analysis of individual RNA-binding domains (left) and schematic representation of the resulting mutants used for binding assays (right). RRM1 was inactivated by mutation of critical aromatic RNP residues, whereas the KH domains were inactivated by GxxG to GDDG conversion. (**b**) EMSAs of the IMP3 mutants with the SELEX-derived 101-mer RNA (see **Fig. 3a**). Mutated IMP3 derivatives (0-120 nM) were titrated to a constant concentration of ^32^P-labeled 101-mer RNA (5 nM). Corresponding binding curves for *K*_D_-estimation are shown on the right, with wildtype IMP3 (WT) included for comparison (mean and standard deviation of three experiments).

Inactivation of the KH3-4 tandem domains also reduced affinity approximately 5-fold, and ΔKH1-2 showed the strongest effect with a 7-fold decreased affinity. These still rather mild effects probably reflect the complex contribution of all tandem domains to overall affinity. Only mutation of all four KH domains (ΔKH1-4) led to a near-complete loss of binding activity. However, note that the observed ΔKH1-4 complexes did not enter the gel, arguing for aggregation of ΔKH1-4 (**Fig. 6b**).

Taken together, this mutational analysis provides further evidence for that all tandem RNA-binding domains of IMP3 actively contribute to RNA recognition.

### SELEX-derived IMP3 consensus in endogenous RNAs

Our findings suggest that IMP3 binds to a complex array of multiple sequence elements that extend over more than 100 nts. For further validation, we analyzed our IMP3-iCLIP data from HepG2 cells^27^, to specifically search for RNAs that harbor the proposed array of binding motifs. Two of the best studied IMP3 targets, *IGF2* and *HMGA2* mRNAs, together with the location of the identified SELEX-motif array are depicted in **Fig. 7a**. The SELEX motifs identified correlate with the highest iCLIP-tag clusters found for the respective mRNAs. Furthermore, we had previously identified *ANKRD17* exon 29 as an IMP3 target that is not only spliced in the canonical mRNA, but that is additionally processed into a circular RNA^27^. Analysis of the exon 29 sequence revealed that it also contains the expected SELEX-motif array with a spacing pattern very similar to our rationally designed 101-mer RNA (**Fig. 7a**, bottom).

**Figure 7.**
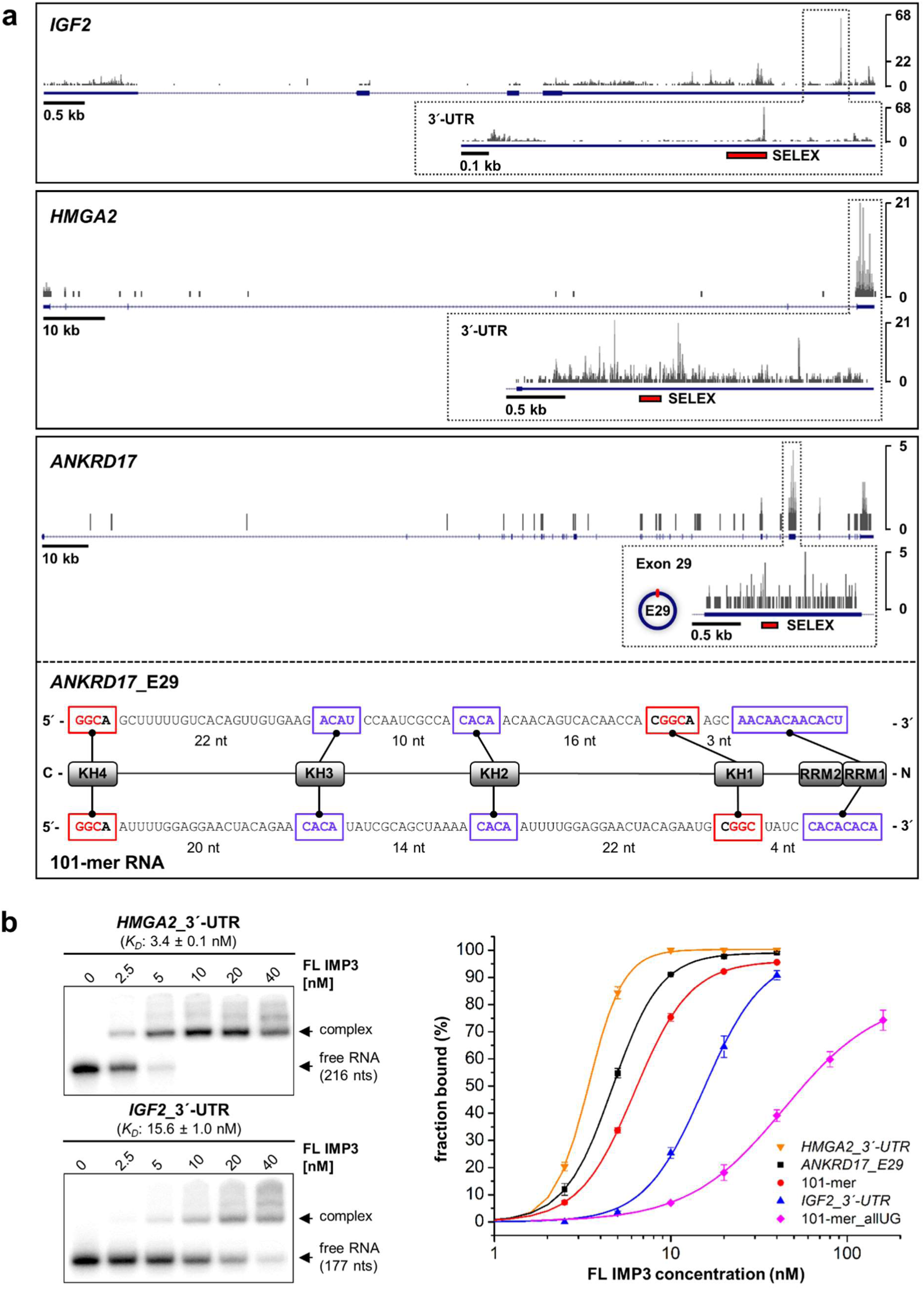
SELEX-derived consensus array in natural IMP3 targets. (**a**) The iCLIP-tag distribution (from HepG2 cells) is schematically represented for three natural IMP3 target mRNAs (*IGF2* and *HMGA2* 3’-UTRs; *ANKRD17* exon 29), with locations of the SELEX-derived motif array indicated by red bars. Exon 29 of *ANKRD17* can additionally be processed into a circular RNA^27^. Below, a detailed schematic of the IMP3 motif array found in *ANKRD17* exon 29 is shown, in comparison to the SELEX-derived 101-mer RNA and including the proposed IMP3-domain-specific recognition of the single RNA elements. (**b**) EMSAs (left) with two natural 3’-UTR targets of IMP3 (*HMGA2* and *IGF2*; 0-40 nM IMP3 and 5 nM ^32^P-labeled RNA). Corresponding binding curves for *K*_D_-estimation are shown on the right in comparison to the SELEX-derived 101-mer RNA and the corresponding negative control 101-mer_allUG (see **Fig. 3b**, mean and standard deviation of three experiments).

Although the isolated RNA sequences from *IGF2* and *HMGA2* 3´-UTRs contain longer spacer regions between the predicted SELEX motifs (exceeding the 25-nt window analyzed), high-affinity binding was measured by quantitative EMSA assays, with *K*_D_ values ranging from 3-15 nM (**Fig. 3b**, for *ANKRD17*, see **Fig. 7b**).

These observations with natural IMP3 target mRNAs further support the biological significance of our SELEX-derived model for RNA recognition of specific sequence elements that can reside in both coding sequences and 3´-UTRs.

### IMP3 regulates the *HMGA2* mRNA by interfering with let-7-mediated repression

Analysis of our iCLIP data had revealed that *HMGA2*, a well-known IMP-regulated mRNA, harbors the IMP3-binding site within a region that also contains two let-7 miRNA seed sequences (**Fig. 8a**, yellow box). As previously reported^9^, a similar, overlapping region is targeted by IMP3, thereby interfering with let-7 dependent *HMGA2* mRNA destabilization. To functionally corroborate our analysis of IMP3 RNA-binding characteristics, we inserted this *HMGA2* region (266 nts) into a luciferase reporter construct and measured the effect of IMP3 motif mutations, let-7 seed mutations^11^, and a combination of both on relative luciferase activity (**Fig. 8a**). Respective luciferase reporter constructs were transfected either in standard ES-2 cells (ctr) or in CRISPR/Cas9 genome-engineered IMP3-knockout cells (KO) (**Fig. 8b**).

**Figure 8.**
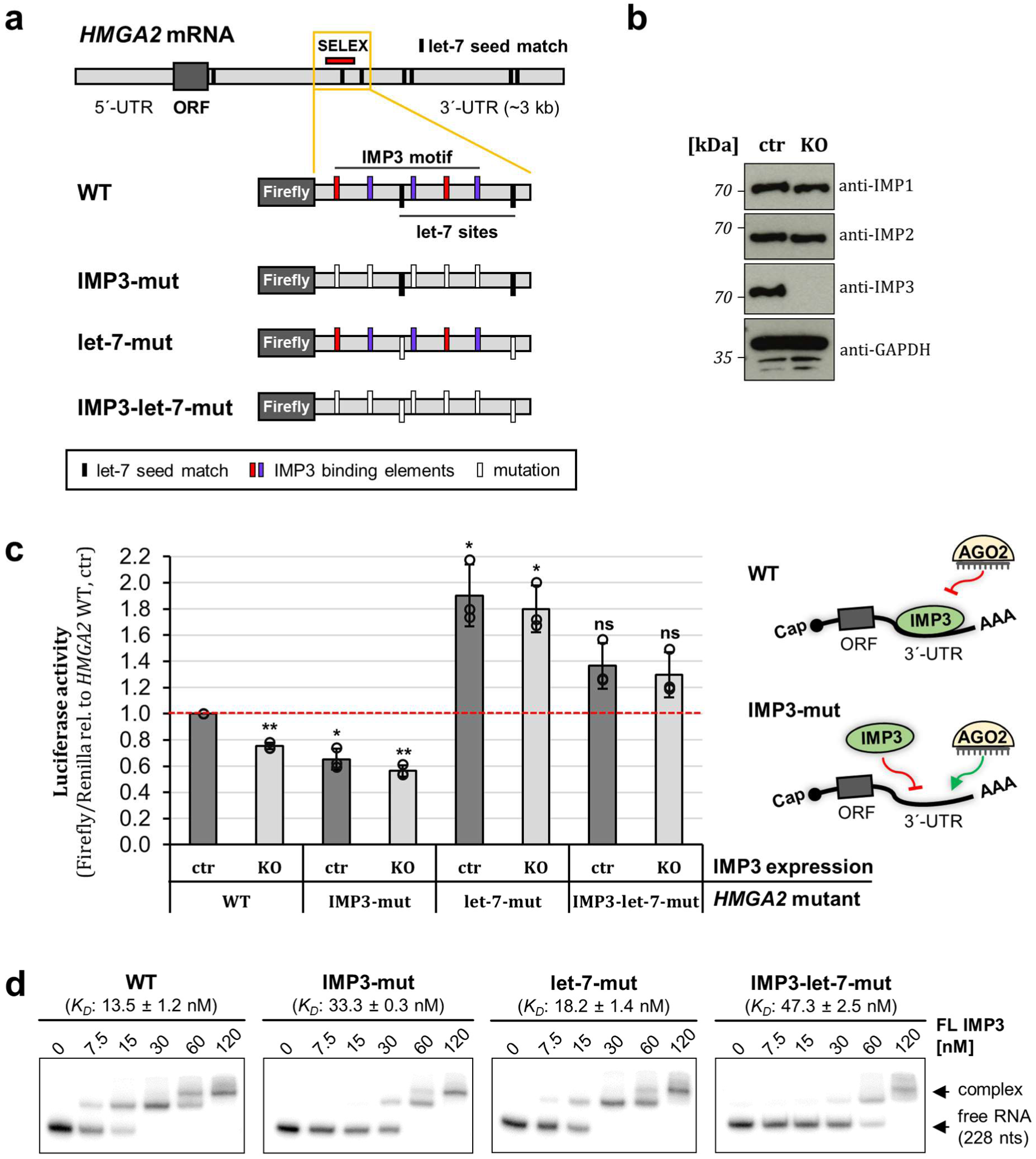
Cross-regulation of *HMGA2* mRNA expression by let-7 and IMP3. (**a**) Schematic of the *HMGA2* mRNA, indicating the seven let-7 miRNA seed matches (black bars) in the 3´-UTR and the SELEX-consensus array (red bar). Below, the structures of luciferase wildtype (WT) and mutant reporters are given, containing the *HMGA2* 3´-UTR region (yellow box) with the IMP3 SELEX-consensus array and two let-7 seed matches. To measure the effect of IMP3 binding, IMP3-binding elements were mutated (IMP3-mut, GGC/CA -> UG, red/violet bars); for analysis of the let-7 influence, the two seed matches in this region were inactivated (let-7-mut, UACCUCA -> UAaCgCA, black bars). In addition, both mutations were combined (IMP3-let-7-mut). (**b**) Western blot analysis of standard (ctr) and CRISPR/Cas9 genome-engineered IMP3-knockout (KO) ES-2 cells, detecting endogenous levels of IMP1, IMP2 and IMP3. GAPDH was used as loading control. (**c**) Standard (ctr) and IMP3-knockout (KO) ES-2 cells were transfected with luciferase constructs described in panel **a**). Luciferase activities were measured as a ratio of Firefly/Renilla activity and compared to control cells transfected with the *HMGA2* WT construct (mean and standard deviation of three experiments; p<0.005^**^, p<0.001^***^, ns = not significant, two-sided t-test). On the right, binding and blocking activities of IMP3 and the let-7-AGO complex within the 3’-UTR of wildtype and IMP3-mutant constructs are schematically represented. (**d**) EMSA assays with ^32^P-labeled *HMGA2* mutant RNAs (0-120 nM IMP3 and 5 nM ^32^P-labeled RNA; mean and standard deviation of three experiments), containing the SELEX motif and a single let-7 seed sequence (see red bar in panel **a**).

In comparison to the WT *HMGA2* sequence, where ∼25% reduction in luciferase activity was observed in IMP3-KO cells, mutation of the IMP3 motif had a more pronounced effect (35% reduction in IMP3-expressing and 45% reduction in IMP3-KO cells), indicating functional inactivation of the IMP3-binding site (**Fig. 8c**). In contrast, mutation of the two let-7 seed sequences increased luciferase activity in both standard and IMP3-KO cells, reflecting the let-7-dependent negative regulatory effect. In addition, by combining both mutations (IMP3-let-7-mut), luciferase activity was slightly, but not significantly increased in comparison to *HMGA2*-WT (WT, ctr), independent of the IMP3 expression status.

To confirm that the observed regulatory effects on *HMGA2* expression are in fact due to changes in IMP3-binding affinity, we performed quantitative EMSAs (**Fig. 8d**). Whereas IMP3 binding to the let-7-mut sequence was nearly unaffected compared to WT *HMGA2*, the affinities for IMP3-mut and IMP3-let-7-mut were decreased 2.5 to 3.5-fold, explaining the activities of our *HMGA2* luciferase constructs.

Taken together, our in-depth analysis of sequence requirements for IMP3-RNA interaction and the functional validation supports the suggested “safe-housing” mechanism: Through sequence-specific formation of RNP complexes, IMP3 shields a specific region within the *HMGA2* 3´-UTR that contains miRNA binding sites in close proximity, thereby protecting the mRNA from let-7-mediated repression.

## Discussion

Members of the IMP protein family are prime examples for multidomain-RBPs, where both affinity and specificity are achieved through simultaneous engagement of multiple domains with their respective RNA elements. Although bioinformatic analyses can predict some features of RNA recognition by multidomain proteins^26,28^ systematic experimental approaches to study combinatorial RNA recognition by multidomain RNA-binding proteins have not been reported so far. Also, commonly employed global approaches to map protein-RNA interactions, such as CLIP, RIP, RNACompete, do not provide such information, but instead, yield only short consensus sequences, thereby severely limiting the systematic description of these RNPs as well as rational searches for high-confidence and functional target sequences^20^.

Here, we focused on IMP3 to dissect its complex RNA-binding through a systematic SELEX-seq approach: We found that all di-domains (RRM1-2, KH1-2, KH3-4) were active in RNA binding while most previous studies had argued that only the KH-domains 3 and 4 guide RNA recognition^5,14-16,29,30^. Our SELEX approach based on N_40_-degenerate sequence revealed that the KH domains recognize two different types of RNA motifs: CA-rich motifs and elements with a common GGC core. Structural analysis of the KH1-2 and RRM1-2 tandem domains and mapping of RNA interactions by NMR unambiguously revealed the specific interaction between subdomains and SELEX-derived RNA motifs. ITC clearly proved a cooperative interaction of tandem KH1-2 with a properly spaced, bipartite RNA motif. Our data suggest that in complex with KH1-2 – similar to the situation with KH3-4 – the RNA adopts a looped conformation that fits the narrow window for linker length between motifs.

In contrast to the IMP1-associated C<u>GGA</u>C motif, we find that IMP3 recognizes two related GGC-core elements (<u>GGC</u>A and C<u>GGC</u>), including their relative arrangement in combination with an additional CA-rich motif. Therefore, our data argue for KH1-2 and KH3-4 acting as independent tandems, both recognizing a combination of one CA-rich motif and one GGC element, with KH1 and KH4 binding the respective GGC elements.

Specifically for KH3-4, and to a lesser extent for KH1-2 and KH1-4, we also observed an enrichment of AU-rich sequences. However, these sequences were underrepresented in full-length IMP3. This may reflect unspecific binding caused by protein truncation. Indeed, C-terminally shortened variants of KH3-4 and KH1-4 were diminished in RNA binding (data not shown).

In contrast to all previous reports^5,29,30^, we found that the N-terminal RRMs also contribute to RNA binding. The analysis of spacing between motifs revealed that all CA-rich motif combinations, but not combinations with the GGC-core elements, were highly enriched in each individual position within the 25 nts window. Most probably, this reflects a specificity for extended CA-repeat elements and binding of several RRM1-2 molecules to CA-rich sequences within the same RNA during the SELEX process. The observed CA-specificity was lost under the stringent washing conditions during SELEX rounds three and four, indicating less robust interactions in comparison to the KH-domains. However, our *in vitro* validation with an RRM1-mutated full-length IMP3 supports an active role of RRM1-2. Based on the conservation of the RNP motifs, we infer that only RRM1 actively contributes to binding, which is supported by our NMR binding data. A model of the RRM1-2 tandem based on NMR and SAXS data suggests that the domains adopt a compact fold, where RRM2 is only indirectly involved in RNA binding, perhaps by stabilizing a compact RRM1-2 arrangement.

Based on these motif analyses with isolated di-domains we designed a prototypic RNA target sequence within a 101-nt RNA that integrates the five SELEX-derived motifs with appropriate spacing. This model was tested and validated by mutational analysis with the 101-mer RNA and *in vitro* binding of well-known IMP3-target mRNAs containing the SELEX-derived motif array (e.g. *ANKRD17, IGF2* and *HMGA2*). In fact, the consensus sequence bound IMP3 with high-affinity, depending on the presence of the individual sequence elements, and involving all tandem RBDs. We observed that isolated tandem domains (e.g. KH3-4) seem to tolerate the enriched motif combinations in both possible arrangements, a phenomenon that was previously described for KH3-4 of IMP1^14,15^. In our spacing analysis, this effect was more pronounced for KH3-4 and KH1-4 in comparison to KH1-2 alone. However, the NMR data of KH1-2 with a corresponding 34-mer RNA ligand indicate a certain degree of dynamic binding judged from the differential line broadening. The dynamic binding could involve the recognition of the 34-mer RNA in both orientations, i.e. with distinct looping of the RNA by the KH tandem domain. Potentially, higher-order oligomers can also be formed at high concentrations of NMR experiments, where for example, line broadening can be caused by the formation of cross-linked RNPs that are in exchange with the 1:1 complex. Interestingly, a clear preference for one orientation (GGC-CA or CA-GGC) was detected for KH1-2 within the full-length IMP3 protein, indicating restricted flexibility of the domains in their canonical context. This is further reflected by a decreased affinity when the order of KH1-2 RNA elements is swapped within the 101-mer RNA. The unique topology may be further enhanced by the kinetic rates of binding, as suggested by Ramos and coworker for looped RNA around KH3-4 at *in vivo* concentrations^16^.

In summary, we provide the first domain-resolved insight into the complex process of IMP3-RNA recognition through concerted interaction of multiple, clustered RNA sequence elements and all RBDs of IMP3. Multivalent interactions of individual domains, each with limited specificity, cooperatively add up to the very specific engagement of full-length protein with target RNAs^22,30^. This greatly exceeds previous studies, including large-scale surveys of many RNA-binding proteins^26,28^, which for the most part were restricted to short recognition sequences. These may even be misleading in many cases, since only particularly dominant sequence elements are usually identified by these approaches. Considering that most RBPs belong to the multidomain type^21,34,35^, our approach presented here on the IMP3 example should advance our understanding of clustered target RNAs^36-39^, and should help in global rational searches for functional target sites as well as in future engineering of tailored multidomain-RBPs^40^.

## Online Methods

### Protein expression and purification

The full-length (FL) and truncated IMP3 derivatives used for SELEX experiments were ordered as codon-optimized DNA fragments encoding FL IMP3 (Met1-Lys579), RRM1-2 (Met1-Asn163), KH1-2 (Pro164-Phe376), KH3-4 (Pro377-Lys579), KH1-4 (Pro164-Lys579) (ThermoFisher), with additional His-tag and TEV-cleavage site, and were cloned into the pGEX-6P2 expression vector (GE Healthcare). For detailed information on purification of the GST-IMP3-TEV-His fusion proteins, see reference 27. IMP3 RNA-binding domain mutants were produced by PCR mutagenesis, using the Q5 Site-Directed Mutagenesis Kit following the manufacturer´s instructions (NEB).

For structural studies, RRM1-2 (Lys2-Asp156) and KH1-2 (Lys192-Ile355) tandem-domain expression constructs were cloned from the human IMP3 full-length protein sequence optimized for expression in *E. coli*. The Δ-versions of KH1-2 were created by restriction-free site-directed mutagenesis. Proteins were expressed as thioredoxin fusion proteins comprising an N-terminal His_6_-tag and a TEV cleavage site between thioredoxin and the gene of interest in the pETTrx1a vector (obtained from Gunter Stier, EMBL Heidelberg). RRM1-2 was expressed by inoculating an LB overnight culture with a clone from a freshly prepared BL21 (DE3) LB culture plate supplemented with 0.35 mg/ml kanamycin. The culture was diluted into the medium of interest and grown to an OD_600_ of approximately 0.8 before induction with 0.5 mM IPTG. Cells were then grown for another four to six hours at 37°C before harvesting. Pellets were resuspended in lysis buffer (50 mM Tris, 300 mM NaCl, 4 mM TCEP, 15 mM imidazole, 1 mg/ml lysozyme, 10 µg/ml DNase I, and protease inhibitors, pH 8.0), incubated on ice for 30 min and sonicated. Cleared lysates were subjected to Ni^2+^-agarose beads. After intensive washing, beads were incubated with 500 µg/l culture of TEV protease in lysis buffer for three hours with gentle shaking at room temperature. Subsequently, the bead supernatant was collected, concentrated and gel-filtrated in 20 mM Bis-Tris, 500 mM NaCl, and 2 mM TCEP, pH 6.5. The respective protein-monomer peak was pooled and salt concentration adjusted to 150 mM. For RRM1-2, we included an additional ion exchange chromatography step to reduce the level of nucleic acid contaminations. This was carried out on a 5/5 MonoS cation exchange column (GE Healthcare), running a gradient from 50-1000 mM sodium chloride in 20 mM Bis-Tris and 2 mM TCEP, pH 6.5. Fractions of intact protein were pooled and dialyzed against the final buffer as before.

### SELEX (systematic evolution of ligands by exponential enrichment)

For detailed information on the SELEX selection steps, see reference 27. Briefly, an RNA pool with a degenerated sequence of 40 nucleotides (N_40_) was prepared by T7 transcription. 40 pmol of GST-IMP3 full-length/truncated derivatives or GST alone (as negative control) were used for four rounds of selection with 4 nmol of SLX-N_40_ transcript. The stringency of washing steps was increased for each round of selection. SELEX selections were carried out with the fusion proteins bound to glutathione-Sepharose (GE Healthcare). RNA aliquots from each round were used for barcoding by reverse transcription with the SLX_RX reverse primers. cDNA libraries were amplified by PCR (12 cycles; SLX_Sol-5xN_fwd and SLX_Sol_rev). The final library pool was subjected to high-throughput sequencing on a MiSeq platform (single-read 150 bp, Illumina). PhiX control library was added to increase sample complexity (Illumina). For primer sequences, see **Supplementary Table 3**.

### IMP3 iCLIP

Sequencing data for the IMP3 iCLIP in HepG2^27^ have been deposited in the Sequence Read Archive (SRA) of NCBI under the accession code SRP139915.

### SELEX-seq data analysis

To identify the enriched binding motifs, sequence reads were first sample-barcode sorted, trimmed by PCR primer sequences on both ends, and further random-barcode filtered to obtain 38-to 40-nt sequence tags of the RNA pools for each sample or round (numbers of sequence tags given in **Supplementary Fig. 1**). The numbers of sequence tags (from each SELEX sample/round) containing either one of the 256, 1024 or 4096 possible tetramer, pentamer or hexamer motifs, respectively, were summarized, and the z-score values were calculated for enrichment of each motif (**Supplementary Table 1**). Each SELEX sample/round was normalized to the corresponding GST SELEX rounds (as negative control and for background correction).

For spacing analysis, sequence tags (round 4 for full-length IMP3, KH1-2, KH3-4, KH1-4; and round 2 for RRM1-2) containing two tetramers with a spacing of 0 to 25 nts were summed up, and the z-score values were assigned. For each of the 65,536 possible combinations of two tetramers, the z-score mean values for spacing of 0 to 25 nts were determined for enrichment ranking. Among the top-500 enriched tetramer-combinations identified for the full-length IMP3 positive control, the following were selected and grouped (see **Supplementary Table 2**):

a) top-10 most enriched combinations of two CA-rich sequences;

b) CA-rich sequence on the 5’ end and <u>GGC</u>A element 3’;

c) <u>GGC</u>A element on the 5’ end and CA-rich sequence 3’;

d) CA-rich sequence on the 5’ end and C<u>GGC</u> element 3’;

e) C<u>GGC</u> element on the 5’ end and CA-rich sequence 3’;

f) two <u>GGC</u>-core elements.

For each group, the z-score mean values for individual positions (0-25 nts) were assigned, and represented as a heat map in **Fig. 2a** (top panel). The motif combinations obtained from b) to e) were subsequently used for spacing analysis of the truncated KH-domain-containing derivatives (KH1-2, KH3-4 and KH1-4; bottom panels). For RRM1-2, and in addition to spacing information of CA-rich sequences from a), motif combinations obtained from b) and d) (5´-GGC-CA-3´), as well as c) and e) (5´-CA-GGC-3´), were combined and presented in a summarized format (middle panel).

### Commercial RNAs

The RRM1-2-related RNAs (CA)_5_, (CA)_3_, (GU)_5_, the KH1-2-related GGC and CA 7-mers, and the 34-mer were obtained from IBA (Göttingen) or Eurofins (Ebersberg). Lyophilized RNAs were dissolved in nuclease-free water, heated to 95°C for five minutes, snap-cooled, aliquoted and stored at −80°C.

### Crystallization, diffraction data collection and processing

The crystallization experiments for IMP3 KH1-Δ2 domain were performed at the X-ray Crystallography Platform at Helmholtz Zentrum München. Initial screening was done at 292 K, using 12 mg/ml of protein with a nanodrop dispenser in sitting-drop 96-well plates and commercial screens. Crystals appeared after 1-2 days with sufficient size for X-ray diffraction experiments. The best data set was collected for a crystal grown in 0.08 M magnesium acetate,

0.05 M sodium cacodylate pH 6.5, 30% w/v polyethylene glycol 4,000 (Hampton Research NATRIX screen). For the X-ray diffraction experiments, the crystals were mounted in a nylon fiber loop and flash-cooled to 100 K in liquid nitrogen. Prior freezing, the crystals were protected with 25% (v/v) ethylene glycol. Diffraction data were collected at 100 K on the PX X06SA beamline (SLS, Villigen). The diffraction data were indexed and integrated using *XDS*^44^ and scaled using *SCALA*^45^. Intensities were converted to structure-factor amplitudes using the program *TRUNCATE*^46^. **Supplementary Table 4** summarizes data collection and processing statistics.

### Structure determination and refinement

The structure of KH1-2 domains was solved by the *Auto-Rickshaw* pipeline^47^. Three-dimensional model of KH1-2 domains of the neuronal splicing factor Nova-1 (PDB-ID: 2ann) ^42^ was used as a search model. For the molecular replacement step followed by several cycles of automated model building and refinement, the *Auto-Rickshaw* pipeline involved the following X-ray crystallography software: *MORDA* (http://www.biomexsolutions.co.uk/morda/), *CCP4*^48^, *SHELXE*^49^, *BUCCANEER*^50^, *RESOLVE*^51^, *REFMAC5*^52^ and *PHENIX*^53^. Model rebuilding was performed in *COOT*^54^. The further refinement was done in *REFMAC5*^44^ using the maximum-likelihood target function. The stereochemical analysis of the final model was done in *PROCHECK*^55^ and *MolProbity*^56^. The final model is characterized by R/R_free_ factors of 23.29 / 29.27% (**Supplementary Table 4**). Atomic coordinates and structure factors have been deposited in the Protein Data Bank under accession code 6GQE.

### NMR spectroscopy

For NMR measurements proteins were expressed in M9 media supplemented with 0.5 mg/ml ^15^N ammonium chloride (titrations and relaxation experiments) and 2 mg/ml ^13^C glucose (triple resonance experiments for backbone assignments). Wildtype KH1-2 has additionally been expressed in 99.5% D_2_O following a previously described protocol^57^. All experiments were performed in 20 mM Bis-Tris, 150 mM NaCl, 2 mM TCEP, 0.02% sodium azide and 5-10% of D_2_O. NMR backbone assignments have been obtained using the following experiments: HNCA, HNcoCA, HNCACB, CBCAcoNH, HNCO, HNcaCO and ^15^N-edited NOESYs. All datasets were acquired from Bruker Avance spectrometers of 600-950 MHz proton frequency equipped with triple-resonance cryo-probes using *Topspin 3*.2. Data were processed with *Topspin* and analyzed using the *CCPNMR Analysis* software package^58^ and *SPARKY*^59^. Sample concentrations were 250-650 µM.

RRM1-2 ^15^N relaxation and hetNOE data were recorded from a 300 µM sample using pseudo-3D experiments with the delays 4, 8, 16, 32, 48, 64, 96, 128, 192, 256, 512 and 1024 ms for T1 and the delays 5, 10, 20, 40, 60, 80, 100, 125, 150, 200 and 300 ms for T2. Peak intensities were fitted and plotted with *Analysis*. T_C_ was calculated based on the ratio of R_1_ and R_2_. NMR titrations of KH1-2 versions and RRM1-2 were performed in samples of 50-100 µM protein by adding the denoted stoichiometries of RNA from a 4 mM stock solution. All NMR experiments were carried out at 25°C. NMR backbone chemical shifts of KH1-2 versions and RRM1-2 will be deposited in the BMRB and available upon publication of the manuscript.

### Static light scattering (SLS)

SLS runs were performed on a Malvern *Omnisec* device with an integrated sample changer and equipped with a semi-analytical SD200 10/300 Superdex column (GE). Samples of RRM1-2 had concentrations as indicated; the used sample volume was 125 µl. Runs were performed in buffers as for NMR, but no D_2_O. UV (260 and 280nm), right-angle-light-scattering and refractive index data were analyzed using the integrated *Omnisec* software and molecular weights determined using a *dn/dc* value of 0.185 for protein. Therefore, peak picking and baseline definition was performed automatically or manually. The system was calibrated with 5 mg/ml bovine serum albumin (MW 66.5 kDa) as a standard.

### Small angle X-ray scattering (SAXS)

SAXS experiments were performed in-house or on beamline BM29 at ESRF, Grenoble, France. Sample concentrations were 1-7 mg/ml. Reference runs in buffers were performed multiple times and used for buffer subtractions. Measurements were carried out as technical triplicates in four to ten frames to enable the exclusion of data in case of radiation damage. Data were processed and analyzed with the ATSAS^60^ package version 2.8 including the plot of paired-distance distribution, P(r), the determination of D_max_ and R_g_ and the calculation of Porod volumes and molecular weights with *DATPOROD*. Theoretical scattering curves derived from the KH1-2 crystal structure or RRM1-2 models were calculated with *Crysol*^31^.

### RRM1-2 modeling

Due to the lack of an experimental structure of RRM1-2, we used SAXS data to filter randomized tandem arrangements. Therefore, RRM1 was modeled based on the IMP2 RRM1 NMR structure (PDB-ID: 2cqh) including residues 1-72. For the RRM2 we used the available structure (PDB-ID: 2e44) and adjusted the domain boundaries to residues 80-156. This fragment was in perfect fit with a CS-Rosetta-based structure based on our backbone NMR data. The linker region 73-79 was kept flexible and the two domains used as an ensemble in 10.000 random starting structures generated with *EOM2*^61^ and fitted against the SAXS scattering curve at the highest concentration. We obtained an ensemble of four structures with populations of 60, 20, and two times 10% that showed a X^2^ fit of 1.335. We chose the highest-populated structure, that also represented the most compact moiety (D_max_ of 61 Å) and used it to include the following restraints: The 7-mer linker (residues 73-79) was rationally probed for possible conformations, i.e. the minimum distance between residues 72 and 80 in a U-turn loop (6 Å), within a α-helix (12 Å) or the maximum distance when arranged in a β-strand (26 Å). The first would have led to steric clashes between RRM1 and 2, and since our secondary chemical shift data did not reveal a clear preference for α-helical or β-strand elements we set the distance to be 16 Å. That allows for sufficient flexibility but would still be in line with a high degree of rigidity (see hetNOE and relaxation data) and fulfill the obtained D_max_ of 54 Å when manually arranging RRM1 and 2. In order to satisfy CSPs we included a maximum distance of 30 Å between residues Val35 (central in RNP2 of RRM1) and Ser127 (RRM2). The latter-despite non-functional RNPs in RRM2-still significantly senses the binding of (CA)_3_ RNA, which would approximately comprise a maximum extension of 30 Å. Finally, the relative twist of RRM1 versus RRM2 around the positively charged inter-domain-linker was limited, given the fact that it senses strong CSPs (see Lys77), indicating it could be arranged along with the RNA. As such, we decided to prevent a cross-brace possibility for linker and RNA and suggest the RNA to bind along the RRM1 β-sheet and the linker thereby indirectly interacting with Ser 127/128. Hence, we put a 15 Å distance between the strongly shifting residue Thr115 and Glu55 to impair the free rotation of domains. All two-domain models were used in the program *Coral*^62^ and fitted against the scattering curves until the crucial parameters D_max_, R_G_ and Porod volume were optimized and the model approximately in line with the CSP plot. The final model showed a X^2^ of 1.9 as given in **Supplementary Fig. 6**. Note that the linker is not part of the model.

### Isothermal titration calorimetry (ITC)

ITC measurements were performed with a MicroCal PEAQ-ITC device (Malvern, United Kingdom) in the NMR buffer. In all experiments, RNA was titrated from a stock of 10-20-fold concentration excess to 20-40 µM protein provided in the reaction cell. In a standard ITC run, we used 19 injections of 2 µl with 150 seconds spacing at room temperature with a 750 rpm stirring speed. Raw data were analyzed with the integrated analysis tool and heat production fitted to a one-site binding model. Where appropriate we performed a buffer subtraction.

### Electrophoretic mobility shift assay (EMSA)

RNAs of the 101-mer series were produced and ^32^P-UTP labeled by T7-transcription from annealed oligo cassettes. SELEX-motif containing regions of *IGF2* (NM_001007139.5), *HMGA2* (NM_003483.4) and *ANKRD17* (NM_032217.4) transcripts were PCR amplified and used for T7-transcription and labeling (sequences given in **Supplementary Table 3**). Binding reactions were performed in binding buffer (10 mM Tris-HCl pH 7.5, 150 mM NaCl, 0.5 mM EDTA, 0.5 mM DTT, 0.1% NP-40, 5% glycerol, supplemented with RNaseOUT, as well as tRNA and BSA as non-specific competitors) containing the purified protein (titration from 0-40, 0-80, 0-120 or 0-160 nM) and the ^32^P-UTP labeled RNA (5 nM) in a final volume of 10 µl. The reaction was first incubated for 30 min at room temperature, and then placed on ice for 5 min. Each sample was supplemented with loading buffer (1x TBE, 0.05% bromophenol blue), and loaded onto a cold native 5% TBE gel (containing 5% glycerol) that had been pre-run for 30 min. Electrophoresis was performed for 50 min with 45 mA at 4 °C. Complexed and free RNA was visualized by the Typhoon FLA 9500 Phosphorimager system (GE Healthcare), and quantified by the ImageQuantTL (GE Healthcare) software. Curve fitting using the Hill equation and *K*_D_ calculation from three independent experiments was performed with OriginPro (OriginLab).

### IMP3 pulldown with biotinylated RNAs and Western blot detection

RNAs of the 101-mer series were produced by T7-transcription (T7 High-Yield Kit, NEB) from annealed oligo cassettes and chemically modified by 3´-biotinylation^63^. For pulldown of IMP3 from HeLa cell lysate, 2.5*10^6 cells where lysed in lysis-buffer (50 mM Tris-HCl pH 7.4, 150 mM NaCl, 5 mM EDTA, 1% NP-40, 0.1% SDS) and incubated with 40 pmol of 3´-biotinylated RNA bound to NeutrAvidin agarose beads (ThermoFisher) in a total volume of 200 µl for 30 min at room temperature. Pulldown of recombinant IMP3 was performed by incubation of 10 pmol 3´-biotinylated RNA bound to NeutrAvidin agarose beads (ThermoFisher) with 1 pmol protein in binding buffer (10 mM Tris-HCl pH 7.4, 100 mM KCl, 2.5 mM MgCl_2_ 0.1% Triton X-100) in a total volume of 200 µl for 30 min at room temperature. After three washing steps with washing buffer (1x WB100, 2x WB300 for pulldown from lysate, and 1x WB100, 2x WB600 for recombinant protein; 10 mM Tris-HCl pH 7.4, 100-600 mM KCl, 2.5 mM MgCl_2_ 0.1% Triton X-100), bound protein was released in SDS-sample buffer (50 mM Tris-HCl pH 6.8, 2% SDS, 10% glycerol, 2.5% 2-mercaptoethanol and 0.05% bromophenol blue) and heat denaturation at 95 °C for 10 min. Samples (10% input and 50% pulldown) were analyzed by SDS-polyacrylamide gel electrophoresis (10% polyacrylamide gel) and Western blotting with polyclonal anti-IMP3 antibody (Millipore) against endogenous IMP3, or anti-GST antibody (Pharmacia Biotech) against the recombinant and GST-tagged IMP3 version.

### CRISPR/Cas9 genomic IMP3 knockout

For the CRISPR/Cas9-mediated genomic deletion of IMP3, ES-2 cells were transfected with two CRISPR guide RNAs (psg_RFP_IMP3_1, psg_RFP_IMP3_2) and Cas9 nuclease (pcDNA_Cas9_T2A_GFP), using Lipofectamine2000 (Thermo Fisher) according to manufacturer’s protocol. Single-cell clones were generated by seeding one RFP-and GFP-positive cell per well using flow cytometry (BD FACSAria II). The deletion of IMP3 was validated by Western blotting using paralog-specific anti-IMP3 antibodies (C-terminal clone 6G8, BSBS AB facility; N-terminal RN009P, MBL). CRISPR guide RNAs are summarized in **Supplementary Table 3**.

### Luciferase assays

The IMP3 SELEX-derived motif and let-7 seed sequence containing region from the *HMGA2* 3´-UTR (NM_003483.4), together with respective mutants (IMP3-mut, let-7-mut and IMP3-let-7-mut) were ordered as DNA fragments (**Supplementary Table 3**, ThermoFisher) and cloned into the pmirGLO-Dual-Luciferase miRNA Target Expression Vector (Promega). For luciferase reporter assays, 1.5 × 10^5^ ES-2 cells (with or without genomic IMP3-KO) were seeded per well, in a 12-well plate. Cells were transfected with 250 ng plasmid DNA and 4 µl Turbofect (ThermoFisher), and incubated for 24 h. After three washing steps with 1x PBS (Gibco), cells were lysed in 250 µl 1x Lysis-Juice (PJK) according to the manufacturer’s instructions. Luminescence was monitored for Firefly luciferase, using the Beetle-Juice Kit, and for Renilla luciferase, using the Renilla-Juice Kit (both PJK) with a Centro LB 960 Luminometer (Berthold Technologies). Relative luciferase activities were calculated as a ratio of Firefly and Renilla raw values with three technical replicates per sample and a total of three independent biological replicates. ES-2 cells with and without IMP3-KO were characterized by SDS-polyacrylamide gel electrophoresis (10% polyacrylamide gel) and Western blotting with antibodies specific for IMP1 (clone 6A9, BSBS AB facility), IMP2 (clone 6A12, BSBS AB facility), IMP3 (Millipore) and GAPDH as negative loading control (Sigma).

## Acknowledgements

We thank Rod Snowdon, Christian Obermeier and Stavros Tzigos (Department of Plant Breeding, University of Giessen) for advice and access to their MiSeq sequencer. We acknowledge support by Ralf Stehle from the SAXS facility of the TU Munich. This work was supported by grants from the Deutsche Forschungsgemeinschaft (DFG, SPP 1935, to A.B., D.N., and M.S.) and by the LOEWE program *Medical RNomics* (State of Hessen; to A.B.).

## Author contributions

T.S., S.T. and A.W. carried out cloning, protein expression and SELEX-seq experiments. L.H.H. performed bioinformatic analyses. T.S. and A.W. carried out EMSA assays. T.S. performed functional experiments. S.M. generated genomic knockout cell lines. A.S., M.A. and J.W. carried out cloning and protein expression for structural studies, performed NMR, SLS and ITC. A.S. and M.A. recorded and analyzed SAXS data. M.A., A.S. and R.J. performed crystallization and structure determination. T.S., D.N., S.H., M.S., A.S. and A.B. designed the project and wrote the paper. All authors discussed the results and commented on the manuscript.

